# Rapid prototyping of a multilayer microphysiological system for primary human intestinal epithelial culture

**DOI:** 10.1101/400721

**Authors:** Sanjin Hosic, Marissa L. Puzan, Will Lake, Fanny Zhou, Ryan A. Koppes, David T. Breault, Shashi K. Murthy, Abigail N. Koppes

## Abstract

Here we report benchtop fabrication of multilayer thermoplastic organs-on-chips via laser cut and assembly of double sided adhesives. Biocompatibility was evaluated with Caco-2 cells and primary human intestinal organoids. Chips with Luer fluidic interfaces were economical ($2 per chip) and were fabricated in just hours without use of specialized bonding techniques. Compared with control static Transwell™ cultures, Caco-2 and organoids cultured on chips formed confluent monolayers expressing tight junctions with low permeability. Caco-2 cells on chip differentiated ∼4 times faster compared to controls and produced mucus. To demonstrate the robustness of laser cut and assembly, we fabricated a dual membrane, tri-layer gut chip integrating 2D monolayers, 3D cell culture, and a basal flow chamber. As proof of concept, we co-cultured a human, differentiated monolayer and intact organoids in a chip with multi-layered contacting compartments. The epithelium exhibited 3D tissue structure and organoids formed in close proximity to the adjacent monolayer. The favorable features of thermoplastics, such as low gas and water vapor permeability, in addition to rapid, facile, and economical fabrication of multilayered devices, make laser cut and assembly an ideal fabrication technique for developing organs-on-chips and studying multicellular tissues.

---

Two-dimensional (2D) tissue culture originated one century ago^1^ and remains invaluable for studying biology and developing therapeutics. Nevertheless, 2D cultures inaccurately represent native tissues. In vivo, continuous nutrient supply and waste product removal occurs via luminal flow along with the circulatory and lymphatic systems, which maintain a homeostatic steady state^2^. Native cells experience physical cues such as fluid shear forces^3,4^ or multilateral mechanical stretching^4^ and chemical cues from heterogeneous tissue-tissue interfaces^5,6^. Microfluidic organs-on-chips are cell culture models that recapitulate heterogeneous tissue-tissue interfaces, which integrate continuous media perfusion to maintain biochemical homeostasis and flow-induced shear stress^7,8^.

The predominant embodiment of organs-on-chip is a bi-layer design featuring two channels interfaced by a porous membrane^9-17^ or hydrogel^18,19^. Culturing different cell types on opposing membrane surfaces or in adjacent channels mimics heterogeneous tissue-tissue interfaces. The bi-layer chip has been used to model the blood-brain barrier^17^, the hematopoietic stem cell niche^16^, the gut microbiome-epithelial-immune interface^12,15^, the lung alveolar-capillary interface^10^, and the placental barrier^14^. Future organs-on-chips integrating patient-derived cells may enable personalized medicine^13^. Interconnecting multiple organs-on-chip via an artificial circulatory system, termed body - on - a - chip, maypermitin - vitro pharmacokinetics^20,21^. Despite these advances, organs-on-chips have been concentrated among bioengineering research groups and have yet to transition to mass clinical or diagnostic applications. Chip automation and parallelization remains challenging, and complex, multi-layered (> 2 layers) chips are limited.

Facile, rapid, economic, and reliable organ-on-chip fabrication would promote interdisciplinary adoption and technological development. Organs-on-chips are most frequently fabricated via poly(dimethylsiloxane) (PDMS) soft lithography^10,12,22-24^. The advantages of PDMS organs-on-chips include high feature resolution, biocompatibility, optical transparency, and gas permeability enabling culture oxygenation and pH control in standard CO_2_ incubators.25 But, PDMS organs-on-chips have several drawbacks. PDMS’ gas permeability25 prohibits O_2_ tension control, which is necessary for recapitulation of hypoxic tissue conditions, as seen in the small intestinal lumen^26^. PDMS’ water vapor permeability^25^ results in evaporation induced bubble formation or high osmolarity, which can block flow and impact cell fate and viability^27^. PDMS also absorbs hydrophobic molecules ^25^, complicating drug pharmacokinetic studies. While PDMS easily bonds to both itself and glass via plasma activation, bonding to polymers requires additional processing such as silanization^28^. PDMS soft lithography requires significant microfabrication training and capital infrastructure^9^. Moreover, initial prototyping may require multiple iterations and lithographic mold fabrication can be prohibitively expensive ($150-$500 per design from 3^rd^ party manufacturers). Other investigators have 3D printed microfluidic cell culture models^29,30^, but these single channel devices do not integrate membranes for recapitulating tissue-tissue interfaces. Therefore, in this study, we aimed to fabricate multi-layered, membrane integrated organs-on-chips without PDMS soft lithography.

A single layer epithelium lines the intestinal wall and forms the rate-limiting barrier to drug absorption^31^. Therefore, oral drug absorption in humans can be approximated using an in vitro differentiated, intestinal epithelium^32^. The human colon carcinoma Caco-2 cell line cultured on permeable supports differentiates into a monolayer with some features of the native small intestine^33^. Organ-on-chip technology was used to develop Caco-2 models with greater fidelity to human intestinal structure and function^11,12,23^. Nevertheless, the immortalized Caco-2 cell line has limited genetic similarity to human intestinal epithelium. Recent advances in intestinal biology have enabled primary human cultures containing intestinal stem cells (self-renewal), Paneth cells (antimicrobial peptide secretion), goblet cells (mucus production), enteroendocrine cells (hormone production), and enterocytes (absorption)^34,35^. Primary, three dimensional (3D) organoids are established from biopsy- or resection-derived intestinal stem cells embedded in Matrigel^34^. While intestinal organoids are genetically and phenotypically more closely related to the native epithelium, organoids form closed lumens that complicate intestinal transport studies. To enable luminal access, researchers cultured primary intestinal monolayers on permeable supports^36-38^, but these models failed to emulate the native 3D tissue structure. More recently, primary intestinal monolayers exhibiting crypt-villus like tissue organization were formed on microengineered scaffolds^39^ and organs-on-chips^13^. Thus, in this study, we too aimed to integrate primary intestinal monolayers and organoids on cut and assembled organs-on-chips.

Here, we describe a “cut and assemble” process for manufacturing thermoplastic organs-on-chips. Most importantly, our technique produced multilayer devices with integrated polymeric membranes and Luer fluidic interfaces faster than soft lithography (hours versus days) at minimal cost ($2 per device) without specialized bonding. The resulting biocompatible, thermoplastic chips are water vapor impermeable, thereby eliminating evaporation-induced bubble formation and osmolarity shifts while potentially enabling O_2_ tension control. The cut and assemble manufacturing technique was validated by reengineering a recently described gut-on-a-chip^12,23^ using Caco-2 cells and primary human intestinal organoids. Caco-2 cells and primary organoids cultured in a bi-layer chip formed confluent monolayers expressing tight junctions and low permeability comparable to static Transwell™ controls. Furthermore, Caco-2 cultures on chip differentiated four times faster towards the enterocyte phenotype as compared to controls and produced mucus, corroborating previously published results.^40^ We integrated primary intestinal monolayers and 3D intact organoids in a dual membrane, tri-layer organ chip. Monolayers exhibited 3D tissue structure spanning 10^2^ µm in height and organoids formed typical cystic structures in close proximity to monolayers, potentially enabling paracrine signaling. The rapid, benchtop, fabrication process presented here has great potential to enable microphysiological modeling of multicellular tissues, 3D cell culture, and the study of paracrine signaling.

## Rapid and facile cut and assemble of bilayer organs-on-chips

The cut and assemble manufacturing technique eliminated PDMS elastomer and microfabrication to overcome the aforementioned limitations of soft lithography^25,27,28^. The bi-layer chip presented here (Fig. 1a-c) featured apical and basal channels interfaced via a PC track etched membrane across a 10 mm length and 1 mm width. The channel height was application dependent. In one embodiment, Caco-2 cells were cultured in a 176 µm tall channel. In a second embodiment to enable monolayer seeding with fewer cells, primary human intestinal cells were cultured in a 1.6875 mm tall channel. Both apical and basal channels had independent inlet and outlets for cell seeding and medium perfusion. Each bi-layer chip consisted of 9 discrete layers (Fig. 1a) irreversibly bonded to membrane interfaced channels (Fig. 1b). While the bi-layer chip featured 9 layers, each device was constructed in 4 steps (Fig. 1d) to align and bond 5 components: (1) a top 3/16” acrylic layer that sealed the chip and provided fluidic connections, (2) a thermoplastic sheet flanked by two adhesive tape layers for the apical channel, (3) a track etched PC membrane to interface the basal and apical channels, (4) a thermoplastic sheet flanked by two adhesive tape layers for the basal channel, and (5) a bottom glass coverslip that sealed the chip. Device geometries were designed in a CAD program and transferred to the respective layers using a benchtop laser cutter. Post laser cutting, 10-32 UNF threads were tapped on the top acrylic layer, enabling commercial Luer lock connections. Then, the device was assembled layer by layer (Fig. 1d).

**Figure 1.**
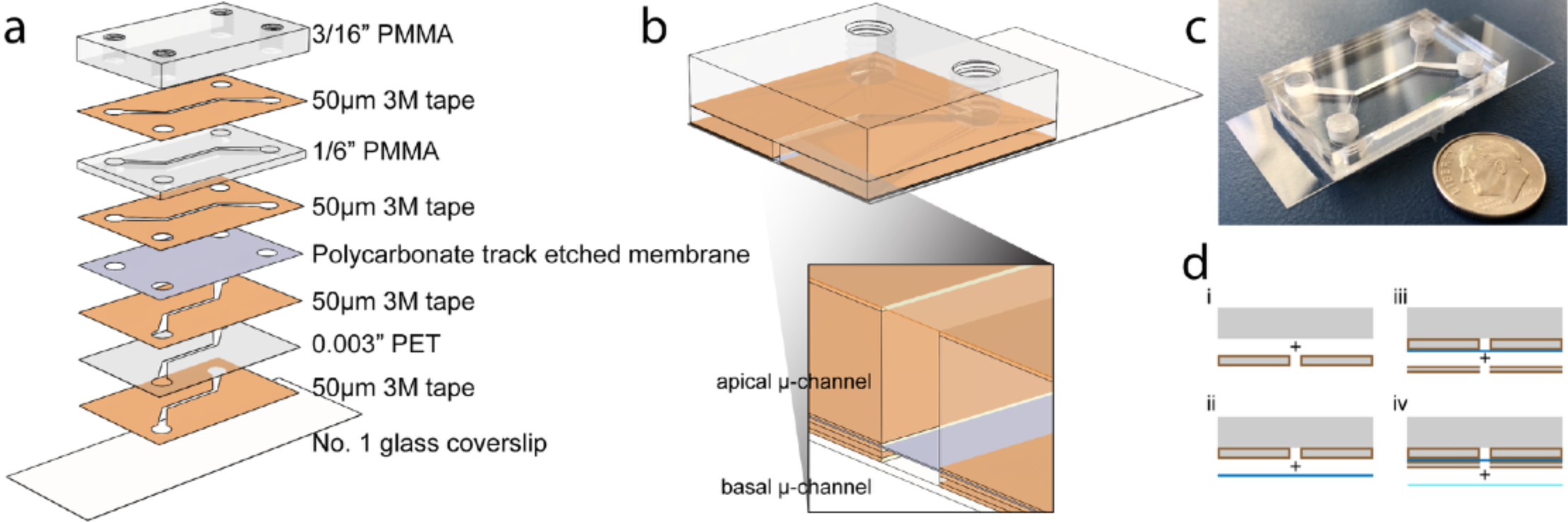
Laser cut and assembled membrane integrated bi-layered organ chip. **a**) A schematic showing the integration of 9 discrete layers to form a bi-layered organ chip. **b**) A schematic of an assembly cutaway view of the bi-layered organ chip showing the inner apical and basal microchannels. **c**) A photograph of the bi-layered organ chip composed of clear PMMA, acrylic adhesive, polycarbonate track etched membrane, and glass coverslip. d) 5 layers are aligned and irreversibly bonded in 4 steps to form the apical and basal channels separated by a polycarbonate track etched membrane with a pore diameter of 1.0 µm.

Cut and assemble fabrication boasts several advantages compared to soft lithography. By eliminating lithographic mold fabrication and the subsequent PDMS degassing, curing, and bonding^9^, chips were cut and assembled in approximately one hour while soft lithography fabrication spans several days^9^. The total material cost per cut and assembled chip was approximately $2. Economical and rapid prototyping is particularly useful in early project phases for iterative design. Eliminating lithography enables researchers without microfabrication experience or facilities to build organs-on-chips. Furthermore, cut and assemble is more amenable to high volume manufacturing techniques such as die cutting, injection molding, and industrial laser cutting as compared to soft lithography.^43^ Though often neglected, facile but reliable fluidic connections simplify chip use, automation, and high-throughput application. Luer Lock and Luer Cone are fluidic interfacing standards but most fabrication techniques, including soft lithography, are incompatible with threaded fluidic connectors. Therefore, we engineered cut and assembled organ chips with threaded ports to accept standard Luer fittings. Cut and assembled organ chips potentially enable O_2_ tension control to recapitulate hypoxic tissues because O_2_ permeability in acrylic is an order of magnitude lower than PDMS. Similarly, PDMS’ high vapor permeability results in significant evaporation relative to the microliter-sized compartments, thereby increasing media osmolarity and compromising cell viability^27^. Evaporation induced osmolarity shifts are a minor concern, however, because most organ chips are operated under perfusion^12,15,22-24,44^.

The channel geometry can be easily modified, and additional channels and membranes can be added. This may be useful when emulating multiple biological barriers such as an epithelium alongside an endothelium^10^. The advantages discussed above are a trade off with reduced feature resolution compared to traditional microfabrication. In the x-y plane, soft lithography reliably produces micron-sized features while laser cutting is limited to 10^2^−10^3^ µm. However, high performance laser cutters claim a focused spot size of 25 µm, thereby approaching lithographic resolutions. In the z plane, traditional microfabrication enables tunable feature height by controlled photoresist deposition. Feature heights of 10^2^-10^3^ µm, however, require multiple spin coatings and long subsequent baking steps. The cut and assemble technology feature heights are partially constrained by adhesive tape thickness but can be increased by layering inert thermoplastic materials such as polyester (PET) film between two adhesive tapes. For example, the Caco-2 bi-layer chip featured 176 µm tall channels composed of a 0.003” PET film sandwiched by two 50 µm adhesive tapes, while the primary cell bi-layer chip featured a 1.6875 mm tall channel composed of a 1/16” acrylic sheet sandwiched by two 50 µm adhesive tapes.

## Recapitulating the human intestine on cut and assemble chips

To assess biocompatibility, human intestinal Caco-2 cells were cultured in the cut and assemble bi-layer chip under apical and basal medium perfusion. Prior to cell seeding, both channels were coated with rat tail type I collagen to promote cell adhesion. The medium flow rate of 0.84 µL/min delivered a shear stress of 0.015 dyne/cm^2^ across the epithelial monolayer as previous work suggested an intestinal lumen shear stress of 0.002-0.08 dyne/cm^2^ ^23^. Caco-2 cells were cultured for 5 days on chip and compared to cells grown on static Transwell™ inserts for 5 and 21 days. After these time points, the cells were fixed and stained for F-actin, tight junction protein ZO-1, and cell nuclei. Cells formed confluent monolayers with comparable morphology, F-actin expression and tight junctions under all three culture conditions (Fig. 2). Importantly, Caco-2 cells formed confluent monolayers across the integrated membrane demonstrating suitability for studying epithelial barrier function. While laser cut, or cutter plotter processed adhesives were previously used to fabricate microfluidic devices, these were analytical microfluidic devices^45-48^. This study demonstrates that acrylic based adhesives are biocompatible channel elements that support human intestinal epithelial cell culture.

**Figure 2.**
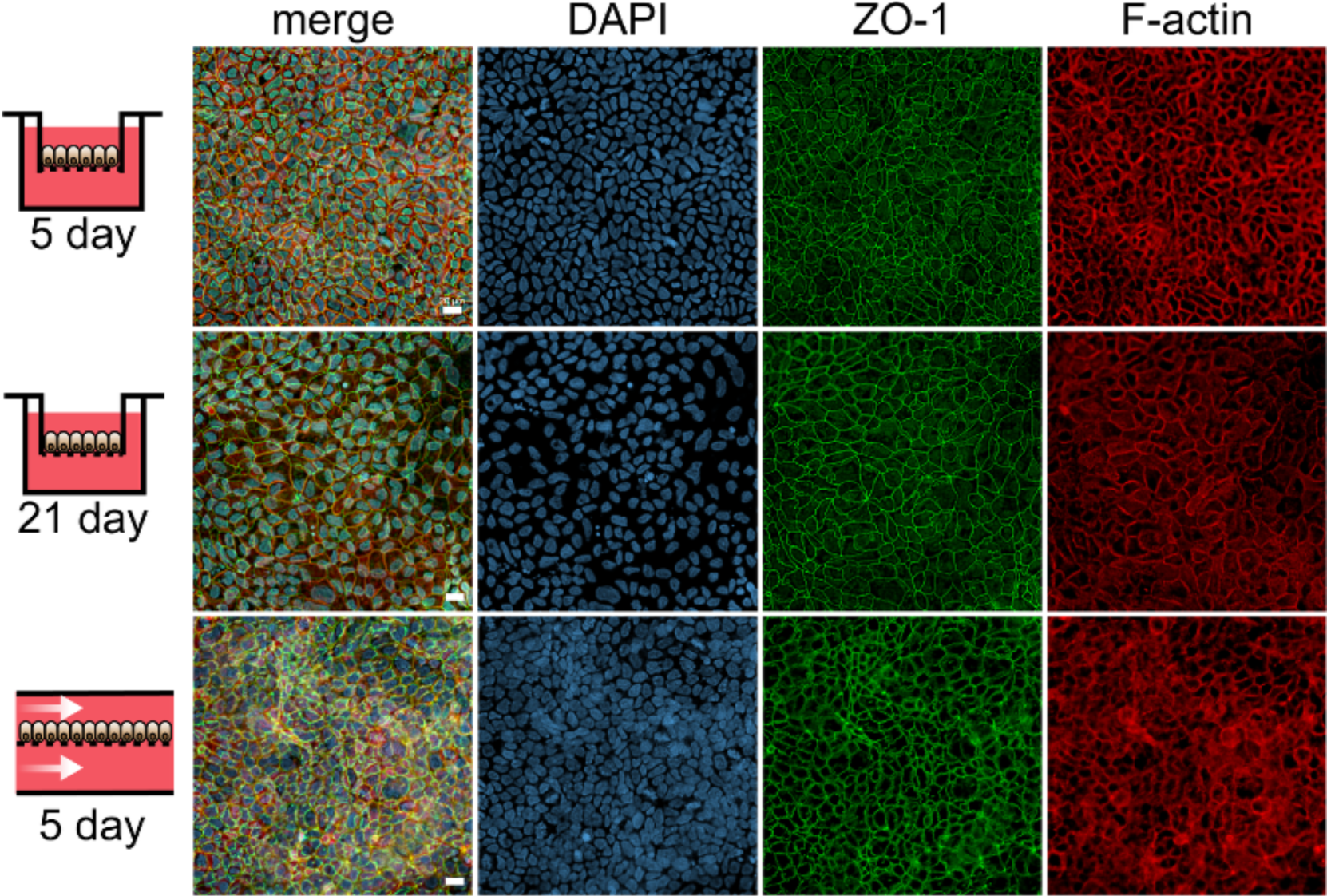
Evaluation of biocompatibility of laser cut and assembled chips. Cell morphology is visualized by immunostaining against ZO-1 tight junctions (green), DAPI nuclei (blue), and F-actin cytoskeleton (red) of Caco-2 cells grown on static Transwell inserts for 5 days (top), 21 days (middle), and on laser cut and assembled chips for 5 days (bottom). Scale bar denotes 20 µm.

## Microfluidic perfusion culture impacts human intestinal function *in vitro*

Following validation of biocompatibility of cut and assemble bi-layer chips we next assessed cellular function. As Caco-2 cells cultured on porous membranes are used to model intestinal transport^49^, we quantified barrier integrity by measuring the apical to basal paracellular transport of a fluorescent dextran (4.4 kDa). Paracellular transport is governed by molecular diffusion through tight junctions rather than active transport via cell membrane bound transporters. The apparent permeability of dextran was calculated as previously described^50^. We did not observe a significant difference (*p* = 0.9748) between the 5 and 21-day static Transwell™ models (Fig. 3a). Furthermore, the results were consistent with previous permeability measurements of a similarly sized dextran across Caco-2 monolayers^51-53^. In contrast, the apparent permeability across the cut and assemble chip Caco-2 monolayers was 40 and 100 times higher than the static 5-day and 21-day Transwell™ models, respectively (*p* < 0.0001) (Fig. 3a). This finding may be partly explained by ongoing perfusion increasing flux at the monolayer surface^54^.

**Figure 3.**
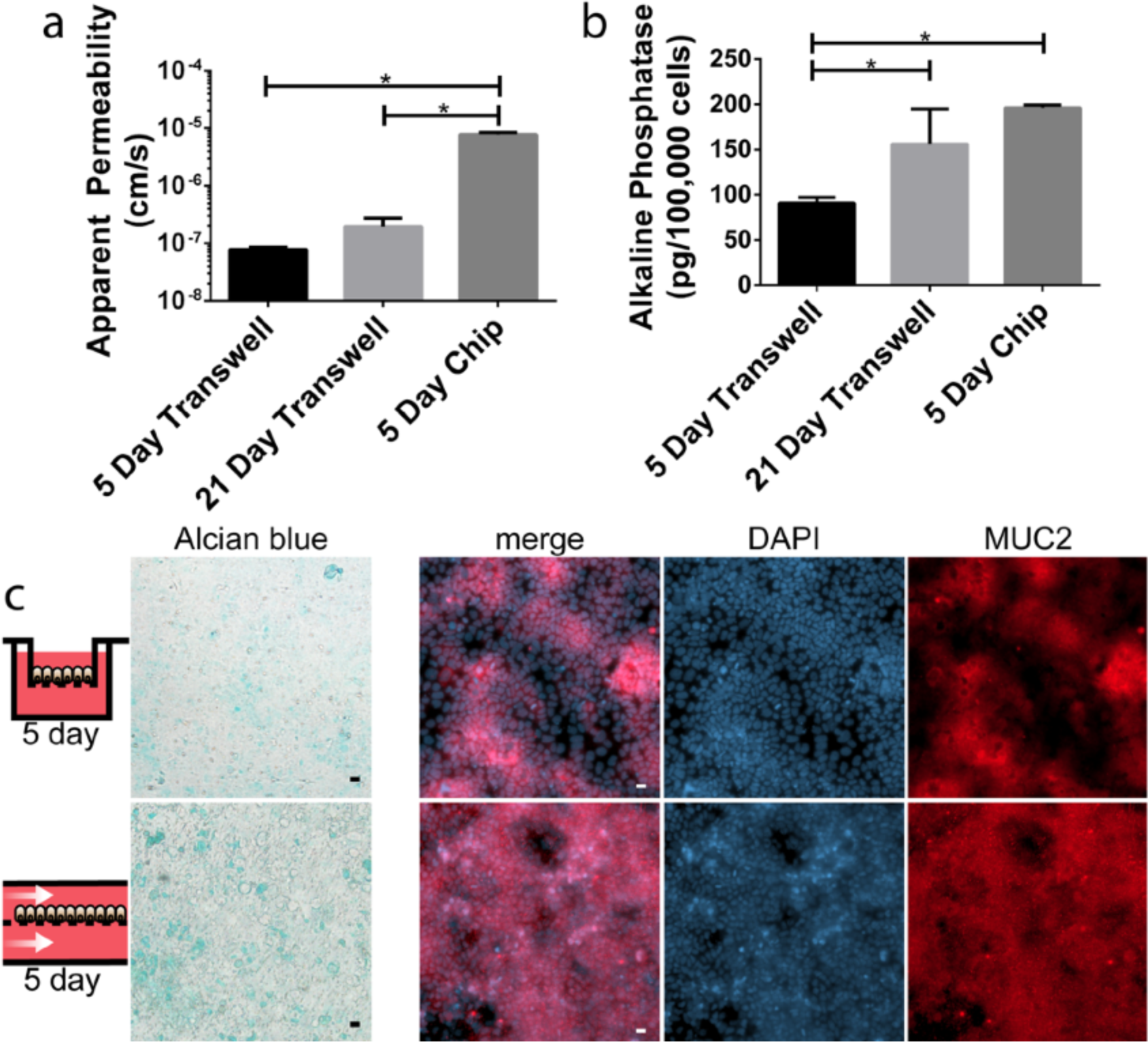
Characterization of epithelial barrier function and Caco-2 monolayer differentiation. **a**) The apparent paracellular permeability quantified by tracking a 4.4 kDa fluorescent dextran through Caco-2 monolayers cultured on Transwell inserts for 5 or 21 days and Caco-2 monolayers cultured on chip for 5 days. Data are presented as means ± STD from 3 independent experiments each utilizing 3 static inserts and 3 chips (* p < 0.05 by ANOVA followed by Tukey’s HSD test). **b**) Alkaline phosphatase expression of Caco-2 cells cultured on Transwell inserts for 5 or 21 days compared to Caco-2 cells cultured on chip for 5 days. Data are presented as mean ± SEM from 3 independent experiments each utilizing 3 static inserts and 3 chips (* p < 0.05 by ANOVA followed by Tukey’s HSD test). **c**) Analysis of mucus production by Caco-2 monolayers on Transwell™ and on chip. Left-most panels depict representative images of Caco-2 monolayers stained with Alcian Blue. In right-most panels, cells are visualized by DAPI nuclei (blue) staining and mucus protein is visualized by anti-MUC2 staining (red) of Caco-2 cells grown on static Transwell inserts for 5 days (top) and on laser cut and assembled chips for 5 days (bottom). Scale bars denote 20 µm.

During extended culture, typically 3 weeks, Caco-2 cells on static Transwell™ inserts differentiate toward an intestinal enterocyte phenotype expressing transport proteins and brush border enzymes^31^. Among brush border enzymes, alkaline phosphatase (AP) is a frequently used differentiation marker^55-57^. As expected, analysis of AP expression revealed a significant 1.7-fold increase (*p* = 0.0317) between Caco-2 on static Transwell™ inserts at 5 versus 21 days (Fig. 3b). However, between Caco-2 cells cultured for = 5 days on chip versus static, AP expression significantly increased 2.2-fold (*p* = 0.0035) (Fig. 3b). The increased AP expression was consistent with a previous study that reported 4-fold increased AP activity by human proximal tubular epithelial cells in response to perfusion.^58^ Perfusion of media may expedite cell differentiation via flow-induced shear stress, increased exposure to nutrients and/or decreased exposure to cellular metabolites and waste products compared to static conditions where media is replenishment every 48 hours.

We next compared mucus production by Caco-2 cells on static Transwell™ inserts versus chip. Alcian blue, a polyvalent dye, was used to identify gastrointestinal mucins^59^ and immunostaining was used to specifically identify Mucin 2, the most abundant structural protein of the gastrointestinal mucus layer^60^. Analysis of alcian blue staining suggested that Caco-2 cells on chip produce more mucus compared cells grown on static Transwells (Fig. 3c). These results are consistent with two previous studies combining Caco-2 and other gastrointestinal cell lines with media perfusion that reported increased mucus production in response to mechanical stimulation via fluid flow^11, 61^.

## Cut and assembled organs-on-chips support primary human intestinal epithelium

While Caco-2 cells on permeable supports are frequently used to model enterocytes for transport studies across the small intestinal epithelium, as a model, Caco-2 are limited by their colorectal adenocarcinoma origin. Caco-2 cells contain unknown genetic mutations, fail to fully recapitulate the gut’s heterogeneous cell population (stem cells, transit-amplifying cells, Paneth cells, goblet cells, enteroendocrine cells and enterocytes), and may not accurately represent any one cell type. Therefore, we sought to establish a more physiologically relevant intestine model by utilizing human primary intestinal epithelial cells expanded as organoids derived from intestinal biopsies (Fig. 4a). Organoids were dissociated primarily to single cells (82%) (Fig. 4b), with 71% viability. Then, cells were plated on either Transwell™ inserts (Fig. 4c) or on cut and assemble chips (Fig. 4d) for 5 or 7 days. Static monolayers were maintained in organoid expansion medium (EM) for 2 days followed by differentiation medium (DM) for 5 days while chip monolayers were maintained in EM for 3 days followed by DM for 2 days. Monolayers were then fixed and stained for F-actin, tight junction protein ZO-1, and nuclei. Primary cells formed confluent monolayers with comparable morphology and tight junctions under both culture conditions (Fig. 5a-b), demonstrating suitability for future epithelial barrier studies. We next quantified barrier integrity (comparing primary monolayers on chip versus on static inserts) by measuring the apical to basal paracellular transport of fluorescent Lucifer Yellow (450 Da). We observed a significant increase (*p=0.0029*) in the apparent permeability between the 5-day static and chip models (Fig. 5c), consistent with observations for Caco-2 cells (Fig. 3a). These results indicate that cut and assemble chips support primary intestinal cells to form confluent monolayers expressing tight junctions and low permeability in response to continuous perfusion.

**Figure 4.**
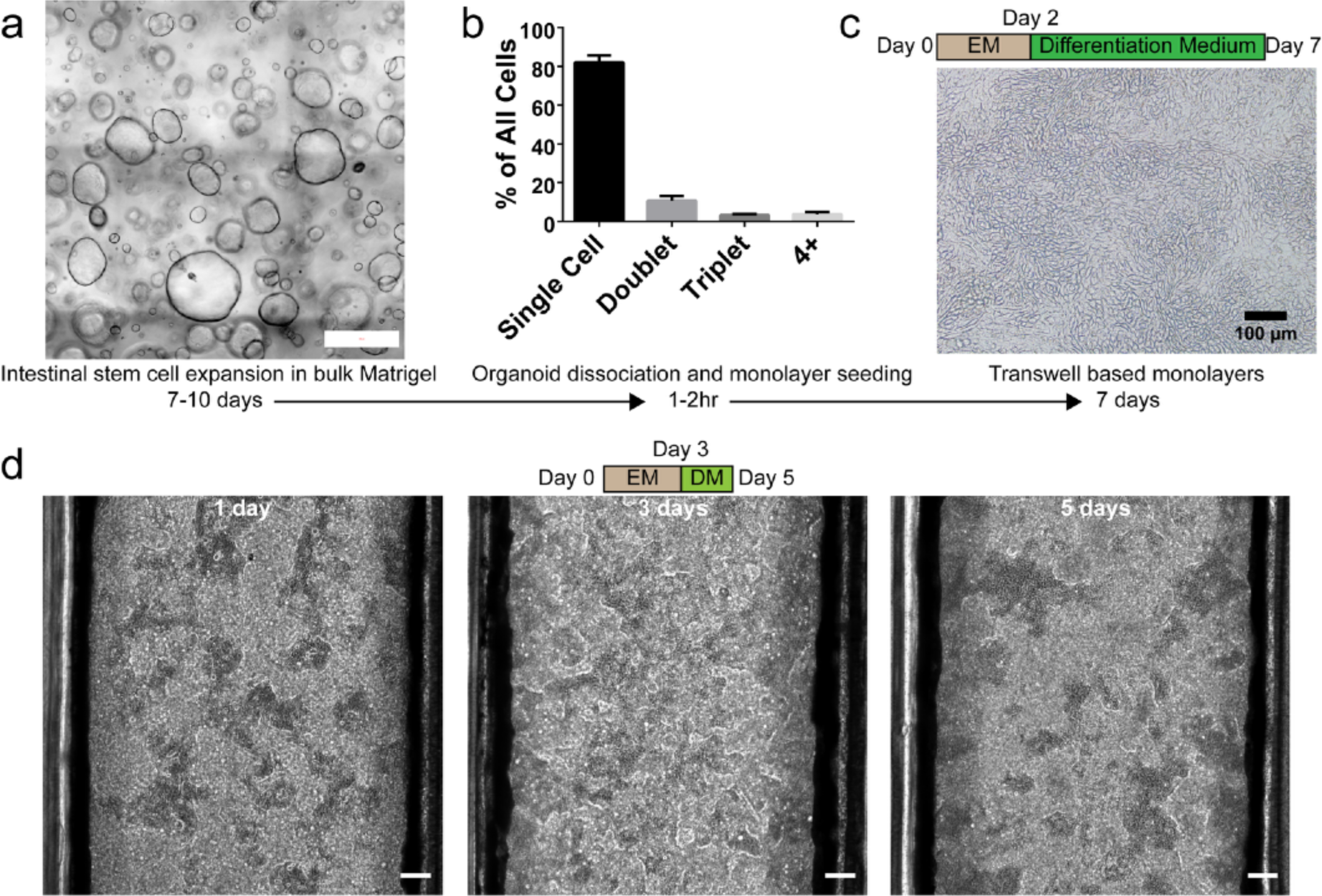
Formation of primary human intestinal monolayers from biopsy derived organoid cultures. **a)** Intestinal stem cells were cultured in Matrigel droplets for 7-10 days at which point the organoids were enzymatically and mechanically dissociated to 82% single cells used for seeding monolayers. Scale bar denotes 500 µm. **b**) the percentage of cells/clumps occurring as single cells, doublets, triplets, or clumps of 4 or more cells after organoid dissociation, n=3. **c**) Transwell™ based monolayers were maintained for 7 days; the phase contrast microscopy image shows Transwell™ primary human epithelial monolayers at 7 days (note that EM was switched to DM at Day 2). **d**) Phase contrast images of primary human epithelial monolayers in the bilayer chip at 1, 3, and 5 days post seeding (note that EM was switched to DM at Day 3). Scale bar denotes 100 µm.

**Figure 5.**
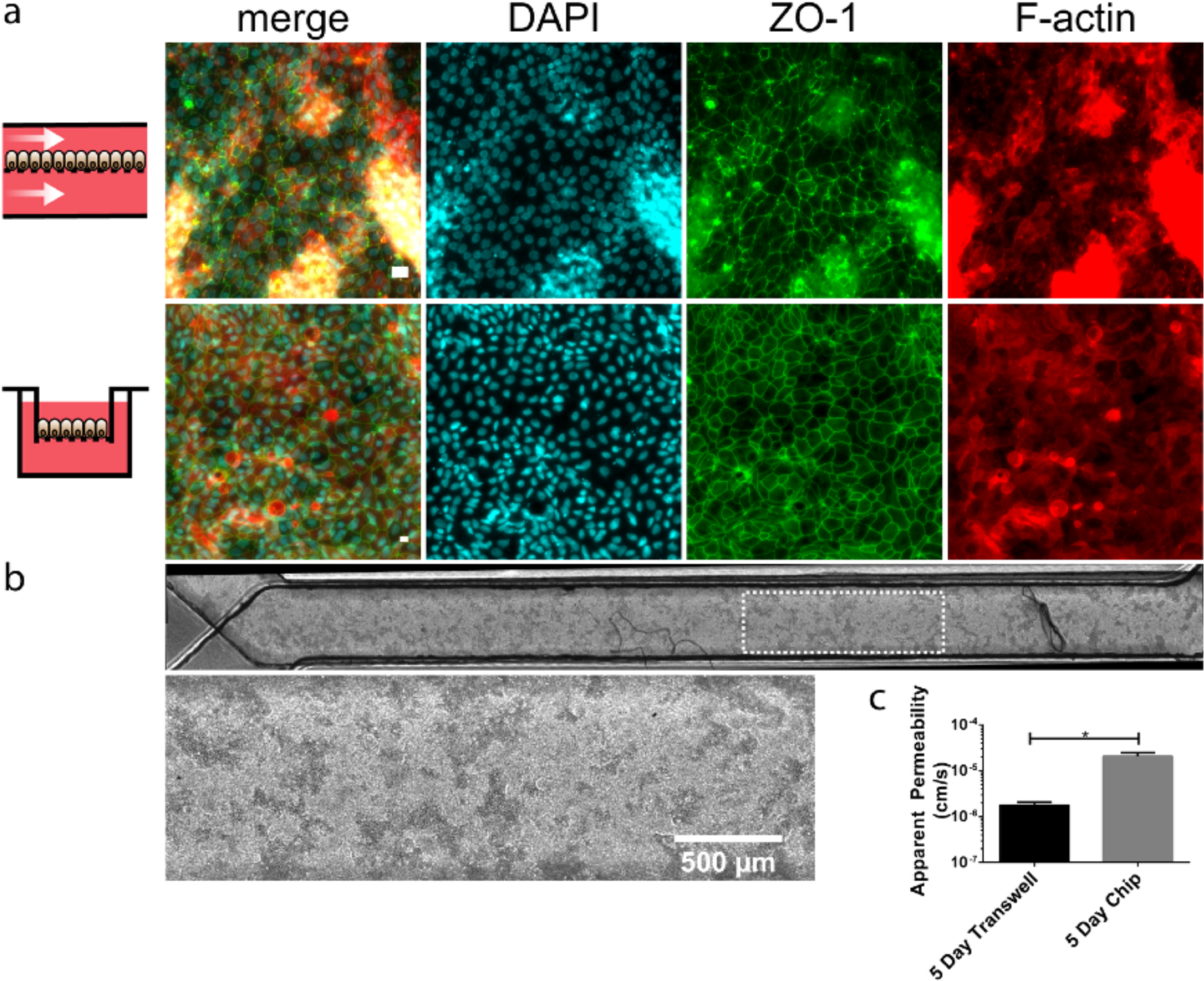
Primary human intestinal epithelium morphology on chip. **a**) Cell morphology is visualized by immunostaining against ZO-1 tight junctions (green), DAPI nuclei (cyan), and F-actin cytoskeleton (red) of primary intestinal cells grown on laser cut and assembled chips for 5 days (top) and static inserts for 7 days (bottom). Scale bar denotes 20 µm. **b**) Phase contrast microscopy shows primary human epithelium at 5 days across the entire length of the chip (top scale bar denotes 1000 µm) and a higher magnification of the area denoted by the white rectangle (bottom, scale bar denotes 500 µm). **c**) The apparent paracellular permeability quantified by tracking Lucifer Yellow through primary epithelial monolayers cultured on Transwell inserts or on bilayer chips for 5 days. Data are presented as means ± STD from 3 independent experiments each utilizing 3 static inserts and 3 chips (* p = 0.0029 by Student’s t-test).

## Integrated primary monolayer and intact organoids in tri-layered organ chips

In the native intestine, epithelial cells are maintained by their surrounding physical, biochemical, and cellular niche^62^. The cellular niche includes myofibroblasts, fibroblasts, endothelial cells, immune cells, glial cells, neural cells, and smooth muscle cells which are embedded in the ECM underlying the epithelium^62^. The cellular niche regulates epithelial cells via both paracrine and contact dependent signaling^62^. While the bi-layer organ chip is the most predominant design, modeling multi-cellular tissues would benefit from more complex organ chip architectures that enable the integration of more than 1-2 cell types or 3D tissue culture within a matrix. To demonstrate the versatility of the cut and assemble method toward fabricating multi-layered architectures enabling 3D tissue culture, we fabricated a dual membrane, tri-layer organ chip (Fig. 6a-c) and integrated 2D and 3D tissue culture of primary intestinal monolayers and intact organoids in adjacent compartments. Organoids cultured in standard plasticware thrive throughout approximately 1.5 mm thick Matrigel (Fig. 4a). Therefore, we engineered the tri-layer organ chip with a 1.6875 mm tall, central, organoid culture channel (Fig. 6c). To promote physical interaction between the differentiated monolayer and the central, organoid laden compartment, the monolayer was cast on a 30 µm pore diameter PC membrane. First, intestinal stem cell laden Matrigel was polymerized in the central channel for 30 min at 37 °C. Then, the apical channel was seeded with dissociated organoids to generate a confluent epithelial monolayer. Organoid expansion medium (EM) was perfused through the apical and basal channels for 6 days. Then, the apical medium was altered to differentiate the epithelial monolayer and perfusion continued for 4 days. Monolayers achieved confluency in 1-2 days and remained confluent over the 10-day culture duration (Fig. 7). Similarly, stem cells formed closed organoids in 1-2 days, which expanded over 10 days and maintained their characteristic cystic morphology (Fig. 7). 3D confocal microscopy of the central organoid channel confirmed the presence of single organoids with their characteristic cystic morphology (Fig. 8c) as well as larger, morphologically complex organoids (Fig. 8d).

**Figure 6.**
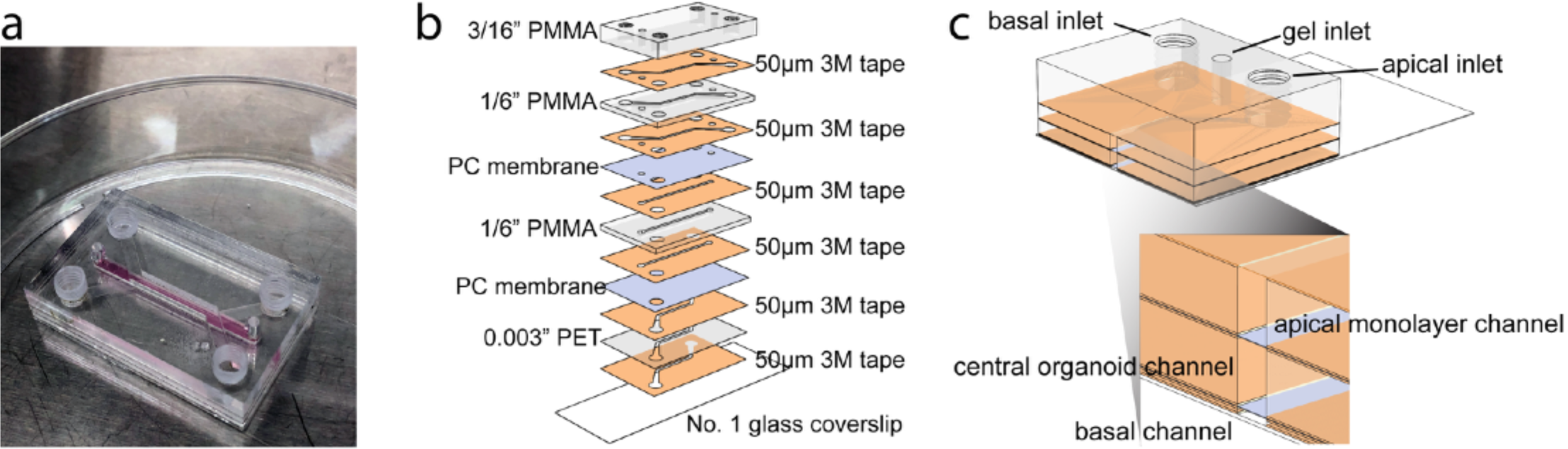
A dual membrane, tri-layered organ chip integrating a primary intestinal monolayer and organoids. **a**) A photograph of the tri-layered organ chip composed of clear PMMA, acrylic adhesive, polycarbonate track etched membranes, and glass coverslip. **b**) A schematic showing the integration of 13 discrete layers to form a tri-layered organ chip. **c**) A schematic of an assembly cutaway view of the tri-layered organ chip showing the inner apical, central, and basal channels.

**Figure 7.**
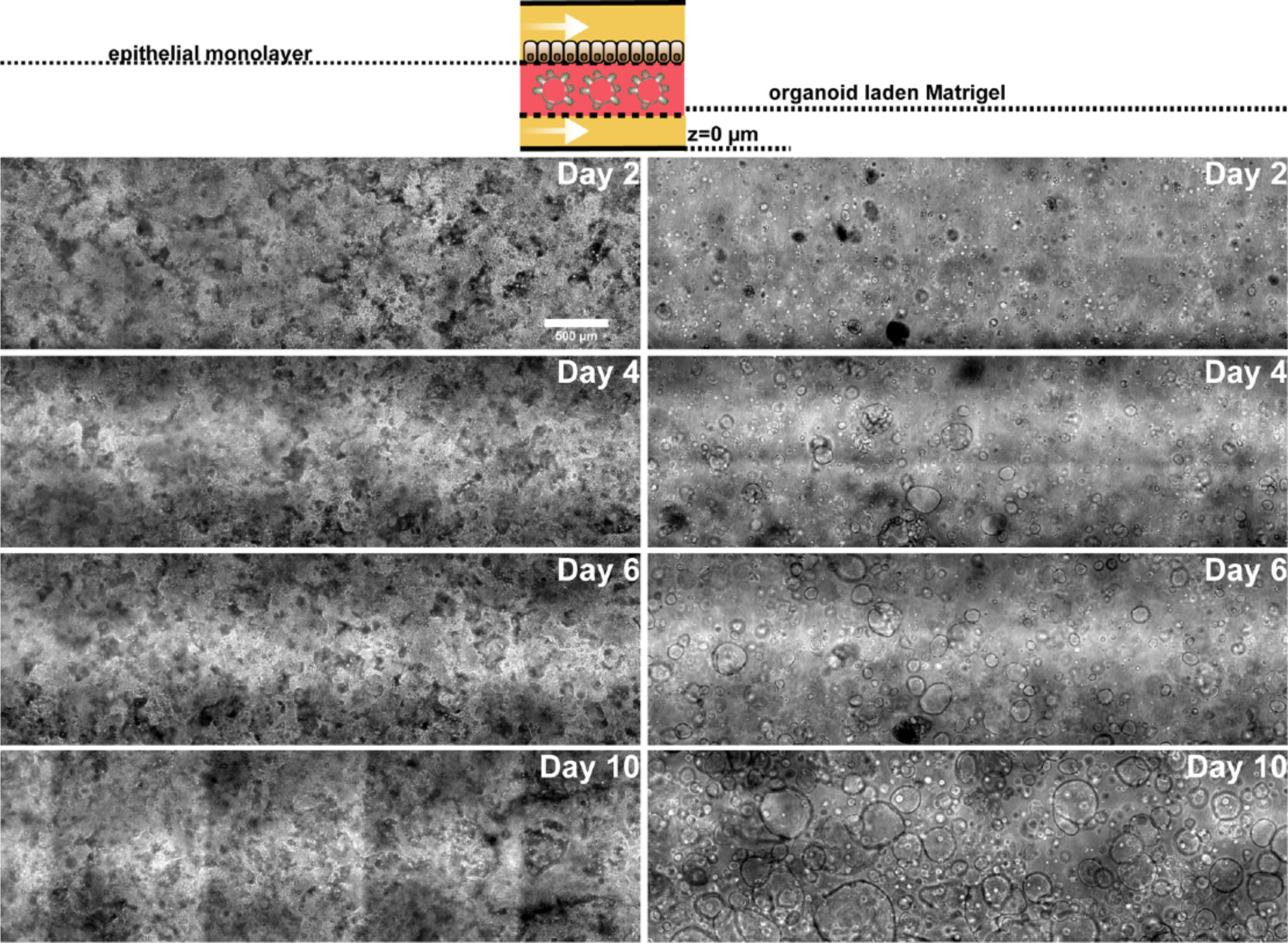
Primary human epithelial monolayers and organoids in the trilayer chip. Phase contrast images at 2, 4, 6, and 10 days post seeding (note that apical Expansion Media was switched to Differentiation Media at Day 6). Scale bar denotes 500 µm.

**Figure 8.**
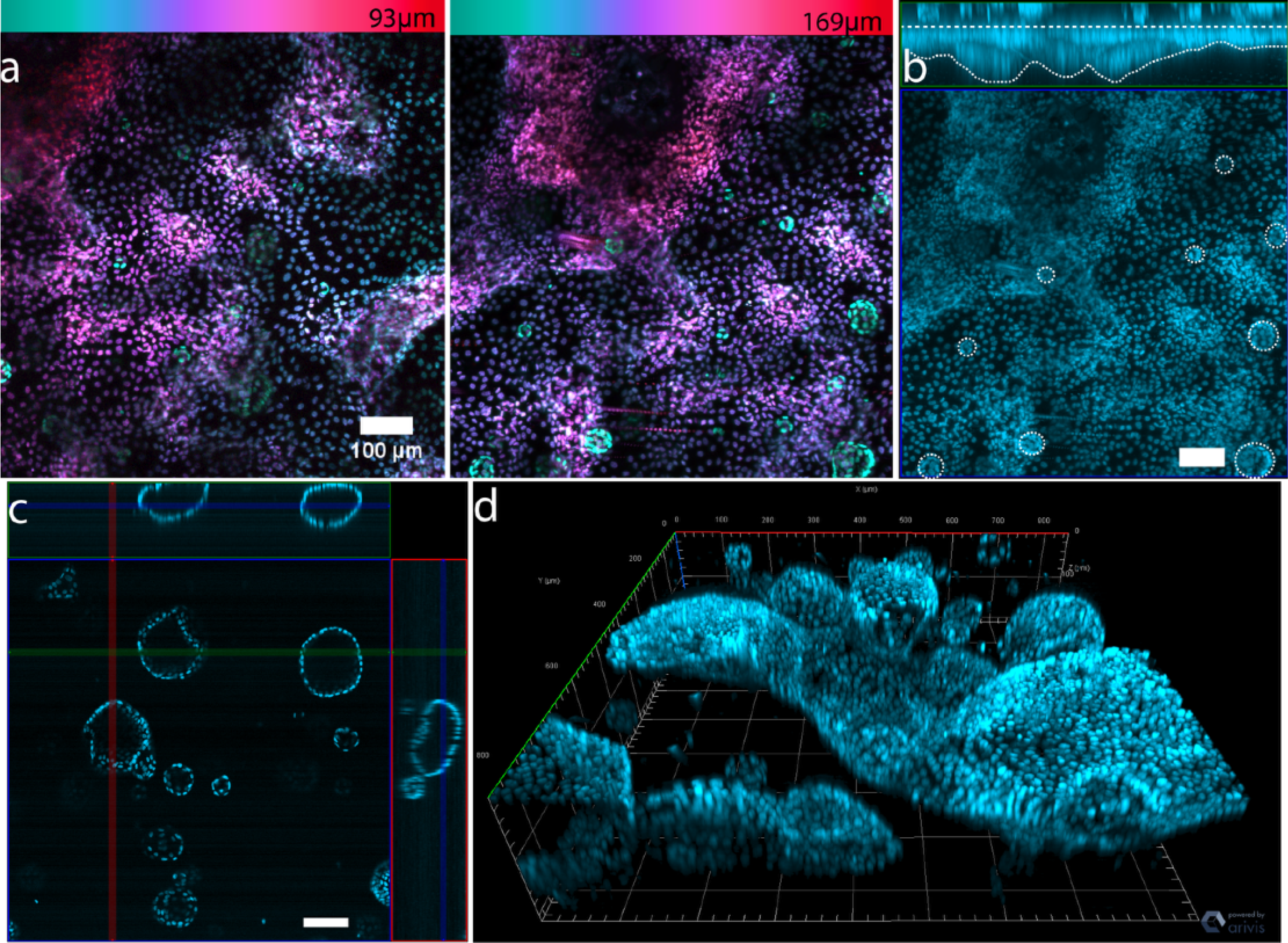
Structural analysis of a dual membrane tri-layered organ chip integrating primary organoid derived monolayers and 3D organoid culture. **a**) Z-depth color coded maximum intensity projections of the monolayer cultured on chip for 10 days and stained with DAPI when viewed from above by confocal microscopy. The color bars above the image specify the range of z-depths in µm. Both images in panel a were observed on the same chip. **b**) A representative maximum intensity z-projection and the corresponding orthogonal view of the monolayer cultured for 10 days and stained with DAPI when viewed from above by confocal microscopy. The dashed white lines indicate the upper and lower surfaces of the primary monolayer while the dashed white circles indicate underlying intact organoids in close proximity to the monolayer. **c**) Representative orthogonal views of intact organoids cultured on chip for 10 days, stained with DAPI, and imaged by confocal microscopy. **d**) Representative 3D reconstruction of confocal immunofluorescent micrographs of intact organoids cultured on chip for 10 days and stained for DAPI. Scale bars denote 100 µm.

While sparse 3D tissue structures spanning a few cell layers were observed in the bi-layer organ chip (Fig. 5a-b), 3D tissue structures in the tri-layer organ chip were present across the entire chip length at an even shorter duration of culture (Fig. 7). To visualize the tissue height, we acquired monochrome 3D confocal stacks and differentially colored each slice to represents the z-depth. Color coded maximum intensity projections of primary monolayers in the tri-layer chip revealed 3D tissue growth spanning approximately 10^2^ µm (Fig. 8a). 3D, multicellular, tissue structures on chip were recently reported and attributed to the presence of media flow and human intestinal microvascular endothelial cells (HIMECs)^13^. Here, we demonstrate 3D tissue growth in the absence of HIMECs. However, the previously reported villus-like structures formed on a collagen I and Matrigel coated membrane. Therefore, it is plausible that biochemical cues originating from laminin and collagen IV rich Matrigel promote 3D tissue growth. In the tri-layer chip presented here tissue growth may be promoted by direct contact between the monolayer and Matrigel through the 30 µm pores as the bi-layer chip was only collagen I coated. Confocal fluorescence microscopy revealed organoids in close proximity to the basal regions of the monolayer and 3D tissue structures (Fig. 8b). It is possible, though unproven, that intact organoids adjacent to the epithelium communicate with the differentiated epithelial monolayer via paracrine signaling to drive morphological changes. For example, intestinal hedgehog signaling in the intervillus pockets of the developing epithelium is involved in crypt-villus axis formation during development and the adult small intestine retains Indian Hedgehog (Ihh) ligands in the differentiated villi^62,63^. Thus, integration of the 2D and 3D microenvironments in the tri-layer gut chip may exhibit more native functionality of intestinal epithelium and stem cells than independent cultures, while allowing evaluation of monolayer and organoid behavior simultaneously. As paracrine Hedgehog signaling between epithelial and mesenchymal cells promotes stromal niche formation which affects epithelial proliferation and differentiation^62^, the tri-layer organ chip presented here is a particularly powerful tool for integrating the small intestine’s mesenchymal components (fibroblasts, endothelial cells, enteric neurons, and glia) and studying paracrine or cell-to-cell contact-dependent (e.g., enteroendocrine cell-enteric glia^64^) signaling.

## Conclusions

Microfabricated organs-on-chips may potentially improve preclinical models, while providing platforms for controlled biological evaluation. In order to commercially succeed, organs-on-chips should be simple to use, automated, and high throughput. Currently, automation and throughput is limited by chip cost and fabrication complexity. The microfabrication free, cut and assemble manufacturing technique presented herein provides rapid (hours), facile, and inexpensive (< $2 per chip) access to multilayer organs-on-chips with standard fluidic connectors. Caco-2 cells cultured on cut an assemble chips formed confluent monolayers expressing tight junctions, which enabled molecular permeability assays. Caco-2 cells cultured on chips also differentiated ∼4 times faster than on Transwell™ inserts, increasing experimental throughput. The cells on chip also produced mucus and alkaline phosphatase, emulating native intestinal functions. Moreover, cut and assemble chips supported primary human intestinal monolayers with tight junction functional barriers.

The versatility of cut an assemble chips was further demonstrated by generating a dual membrane tri-layer gut chip. This tri-layer organ chip may be particularly useful for integrating intestinal monolayers with 3D culture of mesenchymal cells (fibroblasts, endothelial cells, enteric neurons and glial). These cells occupy the ECM rich lamina propria and communicate with the epithelium via paracrine and cell-to-cell contact dependent signaling.^62^ In a proof-of-principle experiment we co-cultured primary 2D monolayers with 3D organoids. Remarkably, the monolayer formed multicellular 3D structures spanning 10^2^ µm, possibly aided by paracrine signaling between the differentiated epithelial monolayer and the proliferative proximate organoids. This platform may enable characterization of intestinal transport and organoid biology towards improved screening and disease modeling, and further design improvements could include enhanced imaging capabilities and increased cell-monolayer interactions. This could be accomplished be decreasing the central channel’s height and/or using an apical membrane with pores > 30µm.

The many features of cut and assemble chips, including the low gas and water vapor permeability of thermoplastics, compared to PDMS, the rapid, easy, and economical fabrication method as well as the ability to make custom multilayered chips, make cut and assemble fabrication well suited for wider adoption and development of organs-on-chips.

## Methods

### Chip design and fabrication

In contrast to practically all previous iterations of gut-on-a-chip, the bi-layer chip discussed herein was fabricated without lithography. The rapid cut and assemble manufacturing process required only a laser cutter/engraver, double-sided adhesive tape, and sheets of poly(methyl methacrylate) (PMMA) and polyester (PET). First, all chip layers were simultaneously designed using CAD software (SOLIDWORKS, Waltham, MA). Then, a laser cutter (Epilog Zing 16, Epilog Laser, Golden, CO) was used to transfer the CAD design to the various materials. The 3/16” PMMA upper layer (McMaster-Carr, Robbinsville, NJ) featured four circular through-holes, which served as fluidic inlets and outlets for both the upper and bottom fluidic layers. The second layer was comprised of a 0.003” polyester or a 1/16” PMMA sheet (McMaster-Carr, Robinsville, NJ), which was sandwiched between 50 µm thick double-sided adhesive tape (966 adhesive, 3M, Maplewood, Minnesota). This 1 mm wide channel served as the upper fluidic channel used for cell culture. The circular inlets and outlets matched the diameter of the through-holes in the uppermost 3/16” PMMA sheet. The third layer, a polycarbonate (PC), track etched membrane with 1.0 µm diameter pores (cat no. 7091-4710, GE Healthcare, Marlborough, MA), featured two circular through-holes with identical diameters as the previous layers. The membrane through-holes enabled fluidic access to the bottom fluidic channel. The fourth layer was comprised of a 0.003” polyester sheet sandwiched between 50 µm double-sided adhesive tape. The features of this layer are identical to the upper fluidic layer but reflected 90°. The fifth and final layer was a No. 1 glass coverslip.

During laser cutting, the protective paper lining on both the PMMA sheets and the double-sided adhesives minimized exposing the materials to burn products. Post laser cutting, 10-32 UNF threads were hand tapped in each of the circular through-holes on the top PMMA sheet. Next, the protective paper lining on both upper and lower PMMA sheets were removed and the plastics were serial cleaned by rinsing with deionized water, Contrad 70 detergent (Fisher), deionized water, isopropyl alcohol and then dried with compressed nitrogen. The laser cut layers were then assembled layer by layer using a custom jig to facilitate hole and channel alignment. The two layers of double sided adhesive tape served as the apical and basal fluidic channels and simultaneously bonded the entire chip (**Figure 1a-c**). Post assembly, the devices were stored under vacuum at 37 °C overnight in order to eliminate outgassing induced bubble formation. Threaded, polypropylene, male Luer lock fittings (cat no. EW-45518-84, Cole-Parmer, Vernon Hills, IL) were connected to the chip via the threaded inlets and outlets. Barbed PC connectors with female luer lock connections (cat no. 11733, Qosina, Ronkonkoma, NY) were connected to the chip. Tubing (cat no. SC-95802-01, silicone, ID 1/32”, OD 3/32”, Cole-Parmer, Vernon Hills, IL) was fitted over the barbed ends to perfuse culture medium through the device.

The bi-layer chip was modified for use with organoid derived primary human intestinal epithelial cells to enable seeding at a lower cell concentration while enabling confluent monolayer formation post-seeding. The top fluidic channel was replaced with a 1/16” PMMA sheet sandwiched between 50 µm thick double-sided adhesive tape (966 adhesive, 3M, Maplewood, Minnesota) for a final channel height of 1.6876 mm. Furthermore, the top 3/16” PMMA cover was not bonded until the day of use at which point the PC membrane was treated with oxygen plasma (50 Watts, 30s, March PX-250 Plasma System). A dual membrane tri-layer chip was designed and fabricated for co-culture of 2D primary monolayers and 3D organoids (**Figure 6a-c**). Two additional components were added to the bi-layer architecture: (1) a 1/16” PMMA sandwiched between two pieces of 50 µm thick double-sided adhesive tape as with the 3D gel channel, and (2) a PC, track etched membrane with 30.0 µm diameter pores (cat no. PCT30047100, Sterlitech, Kent, WA).

### Caco-2 cell culture

Caco-2 epithelial cells were obtained from the American Type Culture Collection (ATCC) and cultured in Dulbecco’s Modified Eagle Medium (DMEM, cat no. 11995-065, ThermoFisher) supplemented with 10% fetal bovine serum (FBS, cat no. 35-011-CV, Corning), and 100 U/mL Penicillin-Streptomycin (cat no. 15140122, ThermoFisher). Cells were cultured in a 37°C, 5% CO_2_ incubator. All experiments were done with Caco-2 cells between passage numbers 40-50.

After fabrication, the bi-layer chip was sterilized via UV irradiation (300 mJ/cm^2^) of the top and bottom chip surfaces (Spectrolinker XL-1000, Spectronics Corporation, Westbury, NY). All tubing and fittings were preassembled and sterilized via autoclave. Both apical and basal fluidic channels were coated with a 400 µg/mL solution of rat tail type I collagen (cat no. 354249, Corning, Corning, NY) in DMEM for at least one hour at 37°C inside a humidified cell culture incubator with 5% CO_2_. The device and tubing were then flushed with Caco-2 culture medium via a sterile, plastic syringe. Caco-2 cells were harvested from a sub confluent T75 flask via 0.25% Trypsin-EDTA (cat no. 25200056, ThermoFisher) and incubation at 37 °C. After cell detachment, the Trypsin-EDTA was diluted with an equal volume of cell culture media and centrifuged at 300g for 5 minutes at room temperature. The cells were suspended in cell culture medium at 5 × 10^6^ cells/mL. The outlet to the bottom fluidic channel was clamped and the harvested cells were infused into the top fluidic channel via a sterile 1 mL syringe. The chips were placed in a 37 °C, 5% CO_2_ cell culture incubator for 1-2 hours for cell attachment. Post attachment, culture medium was perfused through the apical channel via a syringe pump (PhD 2000, Harvard Apparatus, Holliston, MA) at a rate of 0.84 uL/min. The next day, culture medium was perfused through both the apical and basal channels at a rate of 0.84 uL/min.

As controls, Caco-2 cells were cultured on 0.4 µm polyester (cat no. 353095, Corning, Corning, NY) Transwell™ inserts in a 24 well plate. Prior to cell seeding, the inserts were coated with 200 µL of the collagen solution for at least 1 hour at 37 °C inside a humidified cell culture incubator with 5% CO_2_. Caco-2 cells were seeded on the inserts by adding 200 uL of Caco-2 cell suspension (seeding density of 2.6 × 10^5^ cells/cm^2^) and then adding 600 uL of media to the basolateral compartment. The apical and basal cell culture medium was refreshed every other day.

### Organoid culture of biopsy derived intestinal stem cells

De-identified endoscopic tissue biopsies were collected from grossly unaffected (macroscopically normal) areas of the duodenum in children undergoing endoscopy for gastrointestinal complaints. Informed consent and developmentally appropriate assent were obtained at Boston Children’s Hospital from the donors’ guardian and the donor, respectively. All methods were approved and carried out in accordance with the Institutional Review Board of Boston Children’s Hospital (Protocol number IRB-P00000529). Cells were cultured as 3D organoids embedded in 50 µL of Matrigel (cat no. 354230, Corning) on a 24-well plate as previously described^41^. Expansion Medium (EM) for expanding intestinal organoids was prepared from a mixture of Advanced DMEM/F12 medium (cat no. 12634028, Gibco) and 50% L-WRN conditioned medium, which was prepared from L-WRN cells, as previously described^42^. This cell line produces Wnt-3A, R-spondin 3, and noggin. EM was supplemented with GlutaMAX (1x, cat no. 35050061, Gibco), HEPES (10 mM, cat no. 15630080, Gibco), Primocin (0.1 mg/mL, Invivogen), B-27 supplement (0.5x, cat no. 12587010, Gibco), N-2 supplement (0.5x, cat no. 17502048, Gibco), nicotinamide (10 mM, cat no. N0636, Sigma-Aldrich), N-acetyl cysteine (0.5 mM, cat no. A7250, Sigma-Aldrich), epidermal growth factor (50 ng/mL, cat no. 315-09, Peprotech), gastrin (50 nM, cat no. A7250, Sigma-Aldrich), A-83-01 (500 nM, cat no. SML0788, Sigma), prostaglandin E2 (10 nM, cat no. 14010, Cayman Chemical), and SB202190 (10 µM, cat no. S7067, Sigma-Aldrich). Y-27632 ROCK inhibitor (10 µM, cat no. Y0503, Sigma-Aldrich) was added to the organoid medium for the first 48 hours following cell isolation or passage. The culture medium was refreshed every 48 hours using 500 µL per well. Cell culture was performed in a humidified, 37°C, 5% CO_2_ incubator.

Every 7-10 days, the organoids were passaged to new 24-well plates at a ratio of 1:4-1:8 depending on culture density. Matrigel droplets were scratched off the 24-well plate using a 1000 µL pipette tip and collected into a 15 mL conical tube. The organoids were centrifuged at 500g for 5 minutes at room temperature. After aspirating the cell culture medium, the organoids were re-suspended in 0.5 mM ethylenediaminetetraacetic acid (EDTA, cat no. AM9260G, Gibco) in 1x PBS and re-centrifuged at 300g for 5 minutes at room temperature. After aspirating the EDTA, the organoids were re-suspended in Trypsin-EDTA and incubated in a 37°C bath for 2 minutes. The Trypsin-EDTA was then quenched via a 2:1 dilution with Caco-2 culture medium containing 10% FBS and the organoid suspension was triturated ∼10x using a 1000 µL pipette tip to produce single cells and small organoid fragments. The cells were pelleted at 300g for 5 minutes at room temperature. The cells were suspended in 4°C Matrigel and re-plated on new 24-well plates. The plated Matrigel was incubated at 37°C for 15 minutes before adding 500 µL of culture medium containing 10 µM ROCK inhibitor.

### Monolayer culture of primary human intestinal epithelial cells

Polyester Transwell™ inserts in a 24-well plate were coated with 200 µL of collagen solution for at least 1 hour at 37 °C inside a humidified cell culture incubator with 5% CO_2_. Organoids were harvested for dissociation and monolayer seeding after 7-10 days of culture. Matrigel droplets were harvested and processed in Trypsin-EDTA as described above. The Trypsin-EDTA was then quenched via a 2:1 dilution with Caco-2 culture medium containing 10% FBS and the organoid suspension was triturated ∼20x using a 1000 µL pipette tip to produce single cells and small organoid fragments. The cell suspension was filtered through a 40 µm cell strainer (cat no. 22-363-547, Fisher) into a 50 mL conical tube and pelleted at 300g for 5 minutes at room temperature. The cells were resuspended in EM medium with 10 µM ROCK inhibitor. Transwell™ inserts were seeded using 200 µL of cell suspension (seeding density of 9.09 × 10^5^ viable cells/cm^2^) and then 600 µL of media + 10 µM ROCK inhibitor was added to the basolateral compartment. ROCK inhibitor was used for the first 48 hours of cell culture and the apical and basal cell culture medium was refreshed every other day. Following 2 days of culture in EM medium, the apical and basal medium was replaced with differentiation medium (DM) containing Advanced DMEM/F12 + 20% FBS + 4mM GlutaMAX supplement + 100 U/mL Penicillin-Streptomycin. Apical and basal DM medium was replenished every 48 hours.

For seeding primary human intestinal epithelial cells on bi-layer chips, the cells were harvested as described above and suspended at a concentration of 10 × 10^6^ cells/mL. Cell viability was assessed via trypan blue exclusion by incubating cells with an equal volume of 0.4% Trypan Blue Solution (15250061, ThermoFisher). Prior to seeding, the bi-layer chip was treated with O_2_ plasma (50 Watts, 30s, pure O_2_, March PX-250 Plasma System) and the 3/16” acrylic cover was bonded. Note that primary cell adhesion required a plasma treated membrane whereas Caco-2 cells adhered without plasma treatment. The chip was sterilized via UV irradiation as previously described. The chip was coated with collagen solution for 2 hours after which the collagen was flushed with EM medium containing 10 µM ROCK inhibitor. The cell suspension was perfused through the apical channel and the chips were maintained under static conditions in a 37 °C humidified cell culture incubator with 5% CO_2_ for 5-6 hours to enable cell attachment. Then, apical medium was perfused at 1.48 µL/min with EM medium for 3 days before changing to DM medium for 2 additional days. The basal medium was manually refreshed every 24 hours, with EM and DM media, as above.

### Alkaline phosphatase measurement

Alkaline phosphatase (AP) expression was measured using a commercial kit (AS-71109, AnaSpec, Fremont, CA). All kit components were prepared as specified by the manufacturer. Cell lysate from Transwell™ inserts was prepared as follows: medium was removed, and the inserts were washed two times with sterile PBS in both apical and basal compartments. Next, 200 µL of sterile 10x TrypLE^TM^ Select (A1217701, ThermoFisher) was added to the apical side of each insert prior to incubation at 37°C for ∼15 minutes. Cells were collected into a sterile centrifuge tube and inserts were washed with 800 µL of sterile PBS and pelleted at 300g for 5 min at room temperature. The pellet was re-suspended in 150 µL of AP kit supplied buffer, washed and re-suspended in 150 µL of buffer. A 10 µL aliquot of cell suspension was removed to quantify the cell number via hemocytometer. The cells were centrifuged and suspended in 0.2% Triton X-100 (AC327371000, Fisher Scientific). The cells were incubated for 10 minutes at 4°C with agitation and then centrifuged at 2500g for 10 minutes at 4°C. The supernatant was used for the AP assay. 50 µL of the supernatant was moved to a well of a black, polystyrene, 96 well plate (12-566-620, Fisher Scientific). Then, 50 µL of the reaction mixture was added to each well and the plate was manually mixed for 30 seconds. Following a 30 minutes incubation at 37°C, 50 µL of stop solution was added to each well. The plate was manually mixed for 30 seconds. The fluorescence was measured via plate reader (EnSight^TM^, PerkinElmer) using 485nm and 528nm emission and excitation wavelengths, respectively. For Caco-2 on chip, the same protocol was used except that the 10x TrypLE^TM^ and the subsequent PBS wash was infused via a sterile, plastic syringe to detach cells. The actual amount of AP was interpolated using a calibration curve generated with the kit supplied AP standard.

### Paracellular permeability measurement

The apparent permeability coefficient for tetramethylrhodamine (TRITC) labeled dextran (4.4 kDa) (cat no. T1037, Sigma-Aldrich) was determined by measuring transport across the Caco-2 cell monolayer. TEER values of cell monolayers were measured prior to the permeability assay and monolayers with TEER values below 165 Ω × cm^2^ were not used^31^. For control Transwell cultures, 300 µL of 500 µM dextran in cell culture medium was applied to the apical compartment. 100 µL was immediately sampled from the apical compartment, transferred to a black, polystyrene, 96 well plate and stored at 4°C. Transwells were maintained in a humidified, 37°C + 5% CO_2_ incubator. 100 µL aliquots were sampled from the basolateral compartment every 30 minutes over 3 hours and 100 µL of fresh medium preheated to 37°C was added to replace the aliquoted volume. The fluorescence intensity of the collected basolateral samples was measured at 557nm and 576nm emission and excitation wavelengths, respectively. The interpolated dextran concentration was determined using a standard curve. The apparent permeability coefficient was calculated as specified. For Caco-2 on chip, the dextran solution was perfused through the upper channel and cell culture media was perfused through the lower channel at a rate of 0.84 µL/min. Aliquots were sampled from the lower channel every hour and stored at 4°C. The apparent permeability coefficient for Lucifer Yellow (450 Da) (cat no. L0259, Sigma-Aldrich) across primary organoid derived monolayers on static inserts and bi-layer chips was determined as described above.

### Monolayer morphology and mucus production measurement

Cell morphology was assessed by staining F-actin, nuclei, and ZO-1 tight junction protein. Monolayers were washed and stained at room temperature. All monolayers were washed three times with PBS and fixed in 4% formaldehyde (cat no. 28906, ThermoFisher) for 20 minutes. Post fixation, the monolayers were permeabilized in 0.1% Triton X-100 for 20 minutes and blocked overnight in 2% bovine serum albumin solution (BSA, cat no. 97061-416 VWR). The next day, the monolayers were stained with anti-ZO-1 antibody (1:200, 1 hour, cat no. 339188, ThermoFisher), phalloidin (1:500, 1 hour, cat no. A22287, ThermoFisher), and DAPI (1:1000, 10 minutes, cat no. D1306, ThermoFisher) diluted in 1% BSA solution. Transwell membranes were isolated and mounted on a standard glass slide and coverslip with Gold Antifade Mountant (cat no. P36931, ThermoFisher).

Mucus production was assessed by alcian blue and immunostaining for MUC2 protein. Monolayers were washed and stained at room temperature. For alcian blue staining, monolayers were washed three times with PBS and fixed with 4% formaldehyde for 20 minutes. Next, the cell monolayers were washed three times with PBS and stained for 20 minutes with a 1% alcian blue solution in 3% acetic acid (pH=2.5, cat no. 50-319-30, Fisher Scientific) that was filtered via a 0.1µm syringe filter. Post staining, monolayers were washed five times with PBS. For MUC2 immunostaining, the monolayers were washed, fixed, permeabilized, and blocked as previously described. Mucin was detected via an anti-mucin 2 primary antibody (1:200, 1 hour, cat no. PA1-23786, ThermoFisher) and an Alexa Fluor 647 secondary antibody (1:1000, 1 hour, A-21244, ThermoFisher). For monolayers in the bi-layer chip, the protocol remained the same, but washes and stains were applied via syringe pumps to minimize damage to the monolayer.

Fluorescence microscopy was performed on a Zeiss Axio Observer.Z1 microscope equipped with an ORCA-Flash4.0 camera (cat no. C11440-22CU, Hamamatsu). Color images of alcian blue stained monolayers were obtained on an Olympus IX51 microscope equipped with an Olympus DP70 camera.

Confocal microscopy was performed on an LSM 710 confocal microscope (Zeiss) equipped with Zen software (Zeiss) using the Plan-Apochromat 10x/.45 M27 objective. The 405-nm laser was used to excite DAPI. A 512×512 pixel scan format was used. Z-slices were acquired at 2.87-μm intervals with each slice representing the average of 8 scans.

### Statistical analysis

Student’s t-test or ANOVA followed by a Tukey’s post-hoc correction was used to determine statistical significance as indicated in figure legends (error bars indicate standard error of the mean (SEM); p values < 0.05 were considered to be significant).

## Acknowledgements

The authors thank funding support from the National Institute of Health award numbers R21EB025395 Trailblazer (AK and RK) and R01EB021908 BRP (AK and DB) and the Department of Chemical Engineering at Northeastern University for start-up funding (AK and RK).

## Author contributions

S.H., A.K., S.M., conceived the study. S.H., A.K., S.M., and R.K., provided experimental design input. S.H., conducted the experiments, analyzed the data, and was the primary manuscript author. S.H., M.P., W.L., and F.Z., prepared and maintained human organoid cultures. F.Z., and D.B. supplied human intestinal tissue for organoid establishment and WRN conditioned medium. All authors provided input towards the manuscript.

## Additional Information

### Competing financial interests

The authors declare no competing interests.

## References

1 Harrison, R. G., Greenman, M. J., Mall, F. P. & Jackson, C. M. Observations of the living developing nerve fiber. The Anatomical Record 1, 116–128, DOI:10.1002/ar.1090010503 (1907).

2 Li, Z. & Cui, Z. Three-dimensional perfused cell culture. Biotechnol Adv 32, 243–254, DOI:10.1016/j.biotechadv.2013.10.006 (2014).

3 Maggiorani, D. et al. Shear Stress-Induced Alteration of Epithelial Organization in Human Renal Tubular Cells. PLoS One 10, e0131416, DOI: 10.1371/journal.pone.0131416 (2015).

4 Gayer, C. P. & Basson, M. D. The effects of mechanical forces on intestinal physiology and pathology. Cellular signalling 21, 1237–1244, DOI:10.1016/ j.cellsig.2009.02.011 (2009).

5 Powell, D. W. et al. Myofibroblasts. II. Intestinal subepithelial myofibroblasts. Am J Physiol 277, C183–201 (1999).

6 Yoo, B. B. & Mazmanian, S. K. The Enteric Network: Interactions between the Immune and Nervous Systems of the Gut. Immunity 46, 910–926, DOI: 10.1016/j.immuni.2017.05.011.

7 Huh, D., Hamilton, G. A. & Ingber, D. E. From 3D cell culture to organs-on-chips. Trends Cell Biol 21, 745–754, DOI:10.1016/j.tcb.2011.09.005 (2011).

8 Esch, E. W., Bahinski, A. & Huh, D. Organs-on-chips at the frontiers of drug discovery. Nature reviews. Drug discovery 14, 248–260, DOI:10.1038/nrd4539 (2015).

9 Huh, D. et al. Microfabrication of human organs-on-chips. Nat Protoc 8, 2135–2157, DOI:10.1038/nprot.2013.137 (2013).

10 Huh, D. et al. Reconstituting organ-level lung functions on a chip. Science 328, 1662–1668, DOI:10.1126/science.1188302 (2010).

11 Kim, H. J. & Ingber, D. E. Gut-on-a-Chip microenvironment induces human intestinal cells to undergo villus differentiation. Integr Biol (Camb) 5, 1130–1140, DOI:10.1039/c3ib40126j (2013).

12 Kim, H. J., Li, H., Collins, J. J. & Ingber, D. E. Contributions of microbiome and mechanical deformation to intestinal bacterial overgrowth and inflammation in a human gut-on-a-chip. Proc Natl Acad Sci U S A 113, E7–15, DOI:10.1073/pnas.1522193112 (2016).

13 Kasendra, M. et al. Development of a primary human Small Intestine-on-a-Chip using biopsy-derived organoids. Sci Rep 8, 2871, DOI:10.1038/ s41598-018-21201-7 (2018).

14 Lee, J. S. et al. Placenta-on-a-chip: a novel platform to study the biology of the human placenta. J Matern Fetal Neonatal Med 29, 1046–1054, DOI: 10.3109/14767058.2015.1038518 (2016).

15 Shah, P. et al. A microfluidics-based in vitro model of the gastrointestinal human-microbe interface. Nat Commun 7, 11535, DOI:10.1038/ncomms11535 (2016).

16 Torisawa, Y. S. et al. Bone marrow-on-a-chip replicates hematopoietic niche physiology in vitro. Nat Methods 11, 663–669, DOI:10.1038/nmeth.2938 (2014).

17 Sellgren, K. L., Hawkins, B. T. & Grego, S. An optically transparent membrane supports shear stress studies in a three-dimensional microfluidic neurovascular unit model. Biomicrofluidics 9, 061102, DOI:10.1063/1.4935594 (2015).

18 Jeon, J. S. et al. Human 3D vascularized organotypic microfluidic assays to study breast cancer cell extravasation. Proc Natl Acad Sci U S A 112, 214–219, DOI:10.1073/pnas.1417115112 (2015).

19 Adriani, G., Ma, D., Pavesi, A., Kamm, R. D. & Goh, E. L. K. A 3D neurovascular microfluidic model consisting of neurons, astrocytes and cerebral endothelial cells as a blood-brain barrier. Lab on a Chip 17, 448–459, DOI:10.1039/C6LC00638H (2017).

20 Esch, M. B., King, T. L. & Shuler, M. L. The role of body-on-a-chip devices in drug and toxicity studies. Annu Rev Biomed Eng 13, 55–72, DOI:10.1146/ annurev-bioeng-071910-124629 (2011).

21 Edington, C. D. et al. Interconnected Microphysiological Systems for Quantitative Biology and Pharmacology Studies. Sci Rep 8, 4530, DOI:10.1038/s41598-018-22749-0 (2018).

22 Imura, Y., Asano, Y., Sato, K. & Yoshimura, E. A microfluidic system to evaluate intestinal absorption. Anal Sci 25, 1403–1407 (2009).

23 Kim, H. J., Huh, D., Hamilton, G. & Ingber, D. E. Human gut-on-a-chip inhabited by microbial flora that experiences intestinal peristalsis-like motions and flow. Lab on a Chip 12, 2165–2174, DOI:10.1039/C2LC40074J (2012).

24 Jang, K. J. & Suh, K. Y. A multi-layer microfluidic device for efficient culture and analysis of renal tubular cells. Lab Chip 10, 36–42, DOI:10.1039/b907515a (2010).

25 Halldorsson, S., Lucumi, E., Gomez-Sjoberg, R. & Fleming, R. M. Advantages and challenges of microfluidic cell culture in polydimethylsiloxane devices. Biosens Bioelectron 63, 218–231, DOI:10.1016/j.bios.2014.07.029 (2015).

26 Zheng, L., Kelly, C. J. & Colgan, S. P. Physiologic hypoxia and oxygen homeostasis in the healthy intestine. A Review in the Theme: Cellular Responses to Hypoxia. Am J Physiol Cell Physiol 309, C350–360, DOI: 10.1152/ajpcell.00191.2015 (2015).

27 Heo, Y. S. et al. Characterization and Resolution of Evaporation-Mediated Osmolality Shifts That Constrain Microfluidic Cell Culture in Poly(dimethylsiloxane) Devices. Analytical Chemistry 79, 1126–1134, DOI: 10.1021/ac061990v (2007).

28 Aran, K., Sasso, L. A., Kamdar, N. & Zahn, J. D. Irreversible, direct bonding of nanoporous polymer membranes to PDMS or glass microdevices. Lab on a chip 10, 548–552, DOI:10.1039/b924816a (2010).

29 Hamid, Q., Wang, C., Zhao, Y., Snyder, J. & Sun, W. A three-dimensional cell-laden microfluidic chip for in vitro drug metabolism detection. Biofabrication 6, 025008, DOI:10.1088/1758-5082/6/2/025008 (2014).

30 Hamid, Q. et al. Maskless fabrication of cell-laden microfluidic chips with localized surface functionalization for the co-culture of cancer cells. Biofabrication 7, 015012, DOI:10.1088/1758-5090/7/1/015012 (2015).

31 Hubatsch, I., Ragnarsson, E. G. & Artursson, P. Determination of drug permeability and prediction of drug absorption in Caco-2 monolayers. Nat Protoc 2, 2111–2119, DOI:10.1038/nprot.2007.303 (2007).

32 Artursson, P., Palm, K. & Luthman, K. Caco-2 monolayers in experimental and theoretical predictions of drug transport. Adv Drug Deliv Rev 46, 27–43 (2001).

33 Hidalgo, I. J., Raub, T. J. & Borchardt, R. T. Characterization of the human colon carcinoma cell line (Caco-2) as a model system for intestinal epithelial permeability. Gastroenterology 96, 736–749, DOI:10.5555/uri:pii: 0016508589908974.

34 Sato, T. & Clevers, H. Growing Self-Organizing Mini-Guts from a Single Intestinal Stem Cell: Mechanism and Applications. Science 340, 1190–1194 (2013).

35 Leushacke, M. & Barker, N. Ex vivo culture of the intestinal epithelium: strategies and applications. Gut 63, 1345–1354, DOI:10.1136/gutjnl-2014-307204 (2014).

36 Noel, G. et al. A primary human macrophage-enteroid co-culture model to investigate mucosal gut physiology and host-pathogen interactions. Sci Rep 7, 45270, DOI:10.1038/srep45270 (2017).

37 In, J. et al. Enterohemorrhagic Escherichia coli Reduces Mucus and Intermicrovillar Bridges in Human Stem Cell-Derived Colonoids. Cellular and Molecular Gastroenterology and Hepatology 2, 48–62.e43, DOI:10.1016/ j.jcmgh.2015.10.001 (2016).

38 VanDussen, K. L. et al. Development of an enhanced human gastrointestinal epithelial culture system to facilitate patient-based assays. Gut 64, 911–920, DOI:10.1136/gutjnl-2013-306651 (2015).

39 Wang, Y. et al. A microengineered collagen scaffold for generating a polarized crypt-villus architecture of human small intestinal epithelium. Biomaterials 128, 44–55, DOI:10.1016/j.biomaterials.2017.03.005 (2017).

40 Kim, H. J., Huh, D., Hamilton, G. & Ingber, D. E. Human gut-on-a-chip inhabited by microbial flora that experiences intestinal peristalsis-like motions and flow. Lab Chip 12, 2165–2174, DOI:10.1039/c2lc40074j (2012).

41 Sato, T. et al. Single Lgr5 stem cells build crypt–villus structures in vitro without a mesenchymal niche. Nature 459, 262–265, DOI: 10.1038/ nature07935 (2009).

42 Miyoshi, H. & Stappenbeck, T. S. In vitro expansion and genetic modification of gastrointestinal stem cells as organoids. Nature protocols 8, 2471–2482, DOI:10.1038/nprot.2013.153 (2013).

43 Sackmann, E. K., Fulton, A. L. & Beebe, D. J. The present and future role of microfluidics in biomedical research. Nature 507, 181–189, DOI:10.1038/ nature13118 (2014).

44 Phan, D. T. T. et al. A vascularized and perfused organ-on-a-chip platform for large-scale drug screening applications. Lab on a Chip 17, 511–520, DOI: 10.1039/C6LC01422D (2017).

45 Nath, P. et al. Polymerase chain reaction compatibility of adhesive transfer tape based microfluidic platforms. Microsystem Technologies 20, 1187–1193, DOI:10.1007/s00542-013-1901-1 (2014).

46 Sant, H. J. & Gale, B. K. Flexible fabrication, packaging, and detection approach for microscale chromatography systems. Sensors and Actuators B: Chemical 141, 316–321, DOI: 10.1016/j.snb.2009.06.023 (2009).

47 Crews, N., Wittwer, C. & Gale, B. Continuous-flow thermal gradient PCR. Biomed Microdevices 10, 187–195, DOI:10.1007/s10544-007-9124-9 (2008).

48 Kim, J., Surapaneni, R. & Gale, B. K. Rapid prototyping of microfluidic systems using a PDMS/polymer tape composite. Lab on a Chip 9, 1290–1293, DOI:10.1039/B818389A (2009).

49 Stenberg, P., Norinder, U., Luthman, K. & Artursson, P. Experimental and computational screening models for the prediction of intestinal drug absorption. J Med Chem 44, 1927–1937 (2001).

50 Hubatsch, I., Ragnarsson, E. G. E. & Artursson, P. Determination of drug permeability and prediction of drug absorption in Caco-2 monolayers. Nat. Protocols 2, 2111–2119 (2007).

51 Leonard, M., Creed, E., Brayden, D. & Baird, A. W. Evaluation of the Caco-2 monolayer as a model epithelium for iontophoretic transport. Pharm Res 17, 1181–1188 (2000).

52 Buzza, M. S. et al. Membrane-anchored serine protease matriptase regulates epithelial barrier formation and permeability in the intestine. Proc Natl Acad Sci U S A 107, 4200–4205, DOI:10.1073/pnas.0903923107 (2010).

53 Cajnko, M. M. et al. Listeriolysin O Affects the Permeability of Caco-2 Monolayer in a Pore-Dependent and Ca2+-Independent Manner. PLoS One 10, e0130471, DOI:10.1371/journal.pone.0130471 (2015).

54 Karlsson, J. & Artursson, P. A Method for the Determination of Cellular Permeability Coefficients and Aqueous Boundary-Layer Thickness in Monolayers of Intestinal Epithelial (Caco-2) Cells Grown in Permeable Filter Chambers. Int J Pharm 71, 55–64, DOI:Doi 10.1016/0378-5173(91)90067-X (1991).

55 Deng, X., Zhang, G., Shen, C., Yin, J. & Meng, Q. Hollow fiber culture accelerates differentiation of Caco-2 cells. Applied Microbiology and Biotechnology 97, 6943–6955, DOI:10.1007/s00253-013-4975-x (2013).

56 Ferruzza, S., Rossi, C., Scarino, M. L. & Sambuy, Y. A protocol for differentiation of human intestinal Caco-2 cells in asymmetric serum-containing medium. Toxicol In Vitro 26, 1252–1255, DOI:10.1016/j.tiv. 2012.01.008 (2012).

57 Baltes, S., Nau, H. & Lampen, A. All-trans retinoic acid enhances differentiation and influences permeability of intestinal Caco-2 cells under serum-free conditions. Dev Growth Differ 46, 503–514, DOI:10.1111/j. 1440-169x.2004.00765.x (2004).

58 Jang, K. J. et al. Human kidney proximal tubule-on-a-chip for drug transport and nephrotoxicity assessment. Integr Biol (Camb) 5, 1119–1129, DOI:10.1039/ c3ib40049b (2013).

59 Matsuo, K., Ota, H., Akamatsu, T., Sugiyama, A. & Katsuyama, T. Histochemistry of the surface mucous gel layer of the human colon. Gut 40, 782–789 (1997).

60 Schaart, M. W. et al. A novel method to determine small intestinal barrier function in human neonates in vivo. Gut 55, 1366–1367, DOI:10.1136/gut. 2006.096016 (2006).

61 Navabi, N., McGuckin, M. A. & Linden, S. K. Gastrointestinal cell lines form polarized epithelia with an adherent mucus layer when cultured in semi-wet interfaces with mechanical stimulation. PLoS One 8, e68761, DOI:10.1371/ journal.pone.0068761 (2013).

62 Meran, L., Baulies, A. & Li, V. S. W. Intestinal Stem Cell Niche: The Extracellular Matrix and Cellular Components. Stem Cells Int 2017, 7970385, DOI:10.1155/2017/7970385 (2017).

63 van den Brink, G. R. Hedgehog signaling in development and homeostasis of the gastrointestinal tract. Physiol Rev 87, 1343–1375, DOI:10.1152/physrev. 00054.2006 (2007).

64 Bohórquez, D. V. et al. An Enteroendocrine Cell – Enteric Glia Connection Revealed by 3D Electron Microscopy. PLoS ONE 9, e89881, DOI:10.1371/ journal.pone.0089881 (2014).

